# Genomic adaptations in information processing underpin trophic strategy in a whole-ecosystem nutrient enrichment experiment

**DOI:** 10.1101/724484

**Authors:** Jordan G. Okie, Amisha T. Poret-Peterson, Zarraz M.P. Lee, Alexander Richter, Luis D. Alcaraz, Luis E. Eguiarte, Janet L. Siefert, Valeria Souza, Chris L. Dupont, James J. Elser

## Abstract

Several universal genomic traits affect the capacity, cost, and efficiency of biochemical information processing underpinning metabolism and reproduction. We analyzed their role in mediating planktonic microbial community responses to nutrient enrichment in an oligotrophic, phosphorus-deficient pond in Cuatro Ciénegas, Mexico—one of the first whole-ecosystem experiments involving replicated metagenomic assessments. As predicted assuming oligotrophy favors lower information-processing costs whereas copiotrophy favors higher processing rates, mean bacteria genome size was higher in the fertilized treatment, as were GC content, total number of tRNA genes, total number of rRNA genes and codon usage bias in ribosomal protein sequences. However, contrasting changes in trait variances also suggested differences between traits in mediating assembly under oligotrophic versus copiotrophic conditions. Tradeoffs in information-processing traits are apparently sufficiently pronounced to play a role in community assembly and all different components of metabolism— information, energy, and nutrient requirements—are fine-tuned to an organism’s growth and trophic strategy.

## Introduction

Traits influencing the informational underpinnings of metabolism may be crucial to performance and community assembly, but ecologists have largely focused instead on the energetic and stoichiometric features of metabolism (Leal et al., 2017; Sibly et al., 2012; Sterner and Elser, 2002). For an organism to grow and reproduce rapidly, rates at every step of its metabolic network must be sufficiently high such that no single step is unduly rate-limiting, including the information processes that underpin biosynthesis and structure and regulate metabolic networks. This necessary integration of functions is a hallmark of all organisms.

The structure and size of the genome affects the rate, efficiency, and robustness of the information processes supporting metabolism, growth and reproduction (SOM Box 1). There are necessarily tradeoffs in the costs and benefits of these features (SOM Box 1, see also Smith, 1976), which should consequently make individual organisms most competitive and better-suited for only particular ranges of growth and trophic conditions (Roller and Schmidt, 2015). For example, organisms competitive in environments with abundant resources (copiotrophs) must have the capacity for sufficiently high rates of information processing within cells, such as being able to transcribe highly expressed genes and translate resulting transcripts at sufficient rates to maintain and expand protein pools required for achieving rapid growth rates during times of resource abundance. However, maintaining the genomic and structural capacity for rapid growth is costly, potentially placing such taxa at a disadvantage in stable, nutrient-poor environments where growth rates are chronically slow (Giovannoni et al., 2014). Oligotrophic environments may thus instead favor organisms (oligotrophs) that have information processing machinery that is less costly to build, maintain, and operate, thereby increasing resource use efficiency and growth efficiency (Koch, 2001; Roller and Schmidt, 2015). So, there are tradeoffs in the rates and costs of biochemical information processing within cells that can influence the degree to which an organism is optimized for oligotrophy versus copiotrophy. This oligotrophic-copiotrophic strategy continuum is reminiscent of the classic slow-fast life history continuum (Stearns, 1992), especially its lifestyle component (Dobson, 2012; Sibly and Brown, 2007), and classical *r/K* selection theory (MacArthur and Wilson, 2001; Pianka, 1972, 1970) and its developments (Grime and Pierce, 2012; Krause et al., 2014). In conjunction with research on the role of body size and other traits in influencing community assembly (e.g., Fukami et al., 2005; Litchman and Klausmeier, 2008; Okie and Brown, 2009; Roller and Schmidt, 2015), this body of work suggests an important community assembly role for traits associated with biological rates and efficiencies of resource use, growth, and reproduction. It is unclear, however, whether the tradeoffs specifically related to rates and costs of biochemical information processing are sufficiently pronounced to play an important role in evolutionary ecology and the assembly of communities, although there are some promising indications (e.g., Roller et al., 2016).

Here, coupling the use of metagenomics with a trait-based framework that synthesizes theory and hypotheses from genomics, ecology, and evolutionary cell biology, we investigate the role that several universal genomic traits play in determining the response of a planktonic microbial community to nutrient enrichment in a whole-ecosystem experiment. To date relatively few studies have coupled metagenomics with a trait-based framework to clarify the drivers of community assembly (Burke et al., 2011; Chen et al., 2008; Mackelprang et al., 2011; Raes et al., 2011) but even fewer (indeed, none that we are aware of) have deployed such approaches in the context of whole-ecosystem experimentation to test ecologically relevant hypotheses under field conditions. We focus on a set of four information processing traits hypothesized to affect the ability of organisms to obtain the high maximum growth rates (and low minimum generation times) necessary for thriving in copiotrophic environments and/or reduce the energetic and resource requirements necessary to persist under nutrient-depleted conditions:

1. *Multiplicity of genes essential to protein biosynthesis.* Higher numbers of copies of rRNA operons and genes of tRNAs increase overall transcription rates, helping maintain the higher abundances of ribosomes (which are constructed from rRNA and proteins) and tRNAs that facilitate increased rates of protein synthesis (Condon et al., 1995; Roller et al., 2016). However, the higher numbers of genes also come at a cost, as more DNA has to be synthesized, maintained, and regulated, and there may be an increased risk of transcribing unnecessarily large pools of rRNA and tRNA, leading to reduced resource use efficiency and growth efficiency (Roller et al., 2016). A consequence of faster-growing organisms maintaining higher cellular quotas of rRNA is that they should also tend to have higher phosphorus content and low biomass C:P and N:P ratios, since rRNA is phosphorus-rich (Elser et al., 2003, 1996; Godwin et al., 2017; Makino et al., 2003), putting them at a disadvantage under low-P conditions.
2. *Genome size.* Organisms with smaller genomes are predicted to do better in stable and oligotrophic environments, as they require fewer resources (such as P) to maintain and replicate their genomes and can have smaller cells with increased surface-area-to-volume ratios facilitating resource uptake (Giovannini 2015). In contrast, organisms with larger genomes should do better in complex or copiotrophic environments, where they can take advantage of their typically higher intrinsic growth rates (DeLong et al., 2006) and more diverse and flexible gene and metabolic networks (e.g., Konstantinidis and Tiedje, 2004; Maslov et al., 2009) to facilitate substrate catabolism and respond more rapidly to feasts following famines.
3. *GC content – the frequency of nucleotide bases guanine (G) and cytosine (C)*. GC content varies greatly among taxa, but the reasons for this variation are controversial, as multiple different selective and neutral forces may act on the percentage of DNA composed of the nucleotide bases G and C (Bentley and Parkhill, 2004; Hildebrand et al., 2010). However, researchers have proposed that G and C have higher energy costs of production and more limited intracellular availability compared to A and T/U (Rocha and Danchin 2002). Additionally, higher GC content DNA has more nitrogen (Bragg and Hyder, 2004). Thus, lower GC content may be favored in oligotrophic environments. In copiotrophic environments it is less clear what selection, if any, operates on GC content and so GC content in these environments may reflect relaxed selection on GC content.
4. *Codon usage bias*. An amino acid can be encoded by multiple different codons (nucleotide triplets), but these synonymous codons have different kinetic properties, including different probabilities of mistranslation. In highly expressed genes essential to growth (such as ribosomal protein genes), there should be increased selection for biasing the usage of certain synonymous codons over others in order to optimize the accuracy and speed of translation, especially in organisms with fast growth rates (Hershberg and Petrov, 2008; Plotkin and Kudla, 2011; Vieira-Silva and Rocha, 2010).

Further details and background on these traits and our predictions based on them are provided in the Supplementary Online Material.

Various studies have revealed correlations of these genomic traits with performance, generation time, and trophic strategy in a variety of eukaryotic and prokaryotic species and begun to unravel the mechanisms by which these traits influence fitness. Building on this work, it has been shown that some of these traits provide “genomic signatures” of the growth rates of species or communities (Vieira-Silva and Rocha, 2010) and suggested that they may play an important role in ecology (Freilich et al., 2009; Lauro et al., 2009; Vieira-Silva and Rocha, 2010; Weider et al., 2005). However, more work is required to help resolve incongruities in the literature, such as different views on the evolutionary ecology of bacteria genome size (e.g., DeLong et al., 2010; versus Vieira-Silva et al., 2010) (see SOM Box S1). More work is also necessary to develop a less fragmented understanding of the evolutionary and physiological ecology of these genomic traits (Bentley and Parkhill, 2004; Freilich et al., 2009; Vieira-Silva and Rocha, 2010) and their role in community assembly across a wide range of organisms and environments. Existing work also only tends to look at one or two of these traits at a time (Allen et al., 2012; Raes et al., 2007). Importantly, most ecological studies of these traits have been based on studies of microbial isolates, comparative analyses, or sampling across environmental gradients (Allen et al., 2012; Foerstner et al., 2005; Freilich et al., 2009; Raes et al., 2007; Roller et al., 2013; Vieira-Silva and Rocha, 2010). Experimental work examining their role in mediating the structure of communities under natural conditions is extremely limited. Given the complexity of biotic and abiotic interactions shaping communities, that most microbial taxa are uncultivable in isolation (Ho et al., 2017), and that community traits can have widely different responses to temporal/experimental versus geographic variation in abiotic variables (e.g., Sandel et al., 2010), direct experimentation in the field with complex communities is required to better establish the validity of inferences about these purported molecular adaptations for ecological dynamics in nature.

Our study site is Lagunita, an oligotrophic, highly P-deficient pond in Cuatro Ciénegas, Mexico (Lee et al., 2015, 2017). Because of its strong nutrient limitation, this ecosystem offers a useful setting for a fertilization experiment to test these ideas. In this study, we use metagenomic sequencing to evaluate the predictions made above for the role of each trait in governing organismal responses to nutrient enrichment. Our study is noteworthy as one of the first whole-ecosystem experiments involving *experiment-level replicated* metagenomic assessments of community response. If these individual genomic features indeed affect the ability of organisms to survive and reproduce as a function of nutrient availability, then collective measures of these features at the metagenomic level, which aggregates the genomes of all of the individuals constituting a community, should likewise exhibit these characteristics and be reflected in their responses to experimental fertilization (Krause et al., 2014; Wallenstein and Hall, 2012).

## Methods

### Study site description

The whole-ecosystem fertilization experiment took place in Lagunita, a shallow (< 0.33 m) pond roughly ∼12 m x 4 m adjacent to a larger lagoon (Laguna Intermedia) in the Cuatro Ciénegas basin (CCB), an enclosed evaporitic valley in the Chihuahuan desert, Mexico. Despite its aridity, the CCB harbors a variety of groundwater-fed springs, streams, and pools. Past research has also shown that these aquatic environments harbor a high diversity of unique microbiota (Souza et al., 2018, 2006) who have evolved under strong stoichiometric imbalance (high nitrogen (N):phosphorus (P) ratios) and prevalent ecosystem P limitation (Corman et al., 2016; Elser et al., 2005). Lagunita waters are high in conductivity, dominated by Ca^2+^, SO_4_ ^2-^, and CO_3_ ^2-^, have an average molar TN:TP ratio of 122, indicative of strong P limitation, as previously demonstrated in this system during a mesocosm experiment completed in 2011 (Lee et al., 2015, 2017). During the summer season, the pond shrinks substantially and the surface water temperature increases.

### Experimental design

On May 25 2012, prior to initiation of fertilization, five replicate enclosures were established in different parts of the pond; these served as unenriched treatments to serve as reference systems for comparison to the pond after enrichment. Each unenriched mesocosm consisted of a 40-cm diameter clear plastic tube enclosing around 41 L (based on an average depth of 0.33 m at the time mesocosms were installed), a volume that slightly fluctuated during the experiment and decreased very slightly towards the end of the experiment due to evaporation (which decreased the pond volume by 1.4%). The mesocosms were fully open to the atmosphere and sediments. Each mesocosm’s water column was gently mixed periodically during our regular sampling (described below). Thus, with exposure to both the air and the bottom sediments, the unenriched mesocosms were essentially cylindrical “cross-sections” of the ecosystem. The 41-L volume is a typical size for an aquatic mesocosm (e.g., see review of 350 mesocosms by Petersen et al., 1999) and appropriate for microbial studies, encompassing on the order of 30 billion prokaryotic cells (estimated from our cell counts). The pond itself (but not the mesocosms) was fertilized every 3-4 days to maintain PO_4_^3-^ concentration in the water elevated at 1 uM (as KH PO) (SI). We also added NH_4_NO_3_ along with the PO_4_^3^-to achieve and maintain a 16:1 (molar) N:P ratio *in situ* (SI). Fertilizer was added by mixing fertilizer solution with ∼2 L pond water and broadcasting the mixture into all regions of the pond.

We thus performed a sustained whole-ecosystem fertilization with replicate internal unfertilized mesocosms to serve as reference systems. Whole-ecosystem manipulation assures that any experimental responses are ecologically relevant as the system manipulated encompasses the full scale and scope of ecosystem processes that might modulate that response (Carpenter, 1998). Such a whole-ecosystem approach can be especially powerful when coupled to appropriate reference systems. Although our internal unfertilized mesocosms were smaller than the surrounding fertilized pond, we consider them to be pertinent references for investigating the role of genomic traits in community assembly under differing nutrient conditions for several theoretical, empirical, practical, and ethical reasons. While replicate ecosystems for fertilization would be preferred, the availability of multiple ponds at Cuatro Cienegas for such experimentation is extremely limited given the basin’s arid nature. Indeed, true replication of whole-ecosystem manipulations is very rarely achieved. We thus followed the recommendations of Carpenter (1989, 1998): relying on application of a strong experimental treatment applied at the ecosystem scale and informed by previous experimentation (Lee et al., 2017) together with replication of internal reference mesocosms to assess impacts of nutrient fertilization. In this way we maximize the ecological realism of our perturbation by applying it at the ecosystem scale while retaining the ability to compare manipulated dynamics against a benchmark. We prefer a whole-ecosystem fertilization of the pond over fertilizing internal mesocosms within the pond because the smaller enclosures might have provided an artificial view of how microbes respond to a nutrient perturbation at the ecosystem scale.

Given the enormous heterogeneity between communities and water chemistry from different sites within the area, as well as ability of microbes to disperse between ponds, using internal reference systems rather than other whole ponds is arguably more informative as it avoids introducing confounding factors related to variation between ponds (such as contrasting microbial communities) and instead introduces just one potentially confounding factor, the difference in size between the unfertilized mesocosms and the fertilized pond. We thus consider our approach to be the scientifically appropriate one for this area. The design is a natural first step from experiments in small, homogenous bottles or bags. Scaling such experiments across multiple ponds/lakes may be a future step for experimental metagenomic research but not a responsible current step for research in the ecologically sensitive Cuatro Cienegas basin area.

### Field monitoring, sampling, and routine water chemistry

Following initiation of fertilization, the pond and internal unfertilized mesocosms were sampled every 4 days to monitor basic biogeochemical and ecological responses (see SI for water chemistry sampling details). At the end of the experiment (32 days), we sampled for metagenomics: five water samples from the pond itself (fertilized treatment) and one water sample from inside each of the five unfertilized internal mesocosms. It is worth noting that, given the substantial seasonal changes in temperatures and water chemistry, we think comparing metagenomic data of the pond pre-fertilization to the fertilized pond 33 days later would be an inappropriate approach for gaining insight into the effects of fertilization. Thus, we focus on comparing post-fertilization metagenome data with temporally matched data from the unfertilized mesocosms.

The water inside the mesocosm was gently stirred with a dip net prior to sampling. Sampling involved submerging a 1-L polycarbonate beaker just under the surface of the water. Microbes in the water samples were filtered onto sterile GF/F filters (0.7-µm nominal pore size, Whatman, Piscataway, NJ, USA), frozen immediately in liquid nitrogen, and held at <80° C until laboratory DNA extraction, purification, and sequencing. Given the 0.70 µm pore size, extremely small prokaryotes were not part of our metagenomes and so our results do not apply to these picobacterioplankton. If anything, their inclusion would augment predicted community-level trait responses to fertilization, since picoplankton are slow-growers, tend to do poorly in nutrient-rich waters, have small genomes, and so likely decreased in abundance in the fertilized treatment. Routine water chemistry methods were used, as in Lee *et al*. (2015, 2017, see also SI)

### DNA extraction, sequencing, and annotation

DNA was extracted using the MO BIO PowerWater DNA Isolation kit with a slight modification (SI). DNA yield and quality were assessed by PicoGreen assay (SI) and prepared for sequencing on Illumina MiSeq with 12 samples per v2 2×250bp sequencing run. Raw reads were trimmed of barcodes, quality filtered, and rarefied to 100,000 sequences per sample (SI). Two samples from the fertilized treatment and one sample from the unfertilized treatment were left out of subsequent analyses because they had sequencing depths less than 20% of the rest of the samples (whose sequencing depth averaged 2.5 × 10^6^ reads) and low-quality scores. Sequences were phylogenetically annotated using the Automated Phylogenetic Inference System (APIS, SI), which is designed to optimize annotation accuracy (Allen et al., 2012). PCoA was used to visualize a Bray-Curtis distance matrix of the APIS annotations.

### Trait bioinformatics

We used Phylosift (Darling et al., 2014) to annotate non-sub-sampled libraries, allowing for the counting of the number of 16S rRNA Bacteria and Archaea genes in each sample, which averaged 2607. Counting 16S rRNA genes provides a means of evaluating the number of rRNA operons (SI)(Grigoriev et al., 2011). The program tRNAscan-SE v1.4 (Lowe and Eddy, 1997) was used to count the number of tRNA genes in non-sub-sampled libraries (SI). In *SI Bioinformatics and Data Analyses*, we describe our method for ensuring that results were not sensitive to variation in DNA sequencing depth. Bacterial genome sizes were estimated according to methods in Allen et al. (2012)(SI). To quantify the degree of codon usage bias of Bacterial and Archaean ribosomal protein gene sequences, we used the program ENCprime to estimate two commonly used metrics, the effective number of codons (ENC) (Wright, 1990) and a related measure, ENC’ (Novembre, 2002), which additionally accounts for departures in background nucleotide composition from a uniform distribution (Novembre, 2002)(SI).

### Statistics

We used T-tests conservatively assuming unequal variances to evaluate mean differences between treatments. For traits, these tests were one-tailed with the alternative hypothesis based on the predicted differences described in the introduction. General Linear Models with treatment (unfertilized vs fertilized) as a fixed factor were used to determine the amount of variation (R^2^) in traits explained by treatment. Numbers of tRNA genes and rRNA operons were log-transformed before these analyses to improve normality. We also employed single-tailed Poisson rate tests to examine differences in the numbers of rRNA operons and tRNA genes, as the Poisson rate test provides a more powerful test than the T-test for examining differences in the per sequence rate of occurrences of tRNA genes and rRNA operons between treatments (SI). Two-sided Bonett’s tests were used to test for treatment effects on variance between samples of community-level traits. Kolmogorov-Smirnoff tests were conducted to evaluate whether or not within-sample distributions of ENC and ENC’ values of ribosomal protein gene sequences differ between treatments. Finally, we examined how well nutrient enrichment predicted the covariation of these genomic traits along a single axis that quantified molecular adaptiveness to oligotrophic versus copiotrophic conditions. Principal components analysis (PCA) of each samples’ sets of community trait values was used to place each metagenome along a single principal component representing the studied genomic features. Before performing PCA, in order to give equal weight to each trait, variables were first standardized by subtracting means and dividing standard deviations.

We consider a type I error rate of α = 0.10 to provide an appropriate balance between type I and type II error for this study, because this study represents one of the first ecological experiments with metagenomics and, due to conservation considerations and field logistic limitations of time and space within the pond, we had limited replication. In doing so we adopt a modern, balanced approach to the use of frequency statistics in science that aims to avoid over-reliance on significance levels and p-values in judging scientific results (Carver, 1993; Halsey et al., 2015; Rothman, 2016).

We focus on the community response of these genomic traits to varying nutrient conditions, rather than on a detailed natural history of the phylogenetic composition of the community, not only because we are interested in integrating genomic traits with trait-based and microbial ecology but also for pragmatic reasons. There are multiple bioinformatic challenges to precisely resolving the phylogenetic composition of entire microbial prokaryotic communities and ensuring that the phylogenetic biases of various molecular methods do not differ between environments or growth conditions (Kunin et al., 2010; Sunagawa et al., 2013). For instance, there are massive gaps in prokaryotic taxonomic databases (Rinke et al., 2013; Temperton and Giovannoni, 2012), and metagenomes generated from low and high growth communities in oceans have different levels of taxonomy blindness (Kalenitchenko et al., 2018; Sogin et al., 2006; Yooseph et al., 2010). Similarly, DNA sequence assembly introduces a bias, as clonal populations with even low coverage assemble very well while high abundance populations with strain diversity will not assemble well.

## Results

Biomass and chlorophyll *a* concentrations increased substantially in response to nutrient enrichment (biomass: 198% mean increase, P =0.009; chlorophyll *a*: 831% mean increase, P =0.001, Fig. S1), indicating that the pond’s autotrophic biota were nutrient-limited and that our treatment was successful in stimulating growth. The ratio of phosphorus to carbon (P:C) of seston biomass increased with fertilization (P =0.014, Fig 1A), providing additional evidence that organisms in the fertilized treatment had higher growth rates, since high P:C is associated with increased P-limited growth rate due to increased allocation to P-rich RNA required for augmented rates of biosynthesis (Elser et al., 2000).

**Figure 1.**
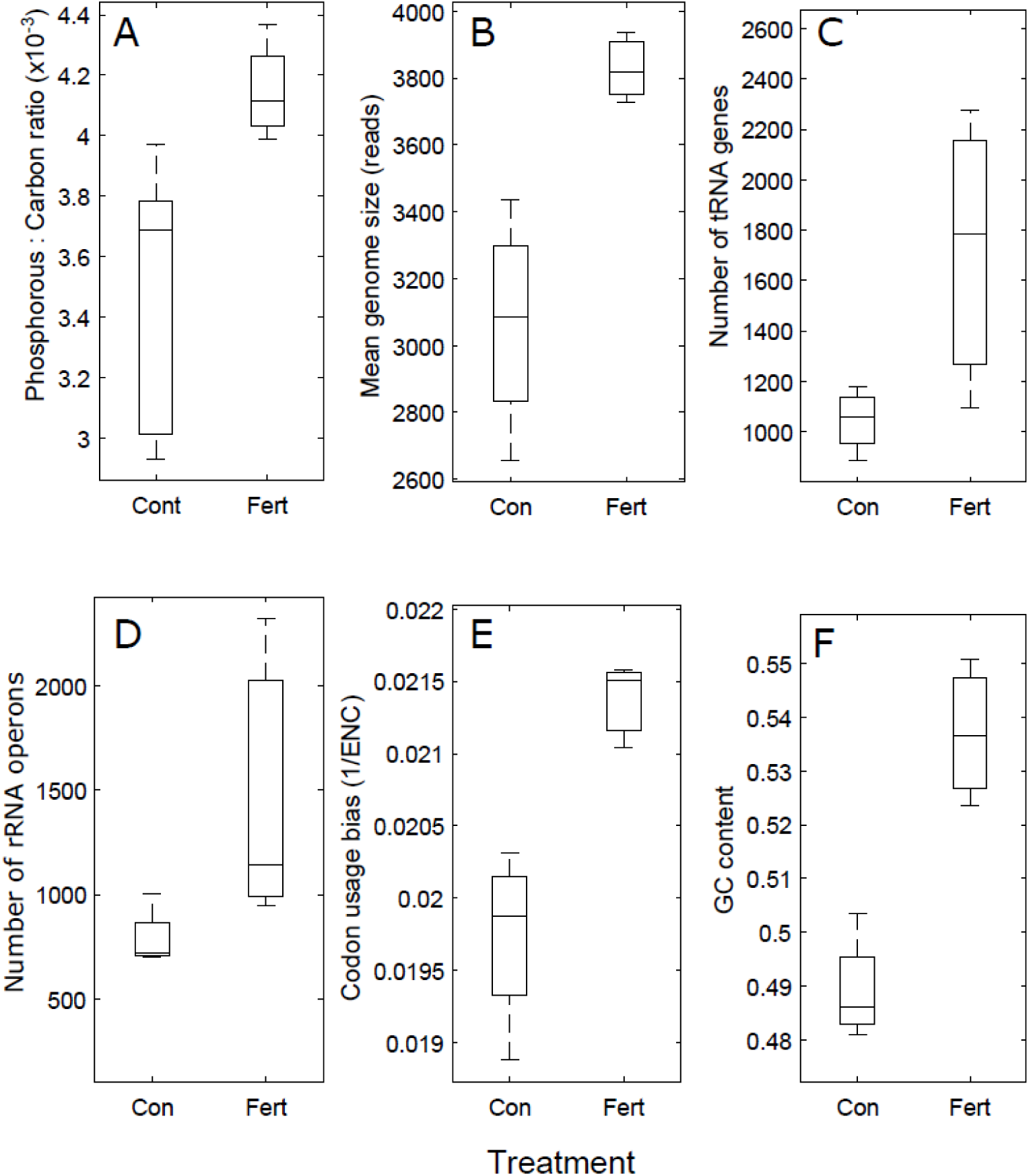
Community-level trait responses to nutrient enrichment. As predicted, each information processing trait was higher in the fertilized treatment (Fert) versus the unfertilized treatment (Con). Because the faster growing organisms generally have more P-rich ribosomes, seston P:C ratio also increased (as shown in **A**). Boxes show 25/75% quantiles, center horizontal lines show medians, and vertical lines show the data range. In **E**, a community’s codon usage bias is the inverse mean effective number of codons (1/ENC) of a metagenome’s ribosomal protein sequences, with higher 1/ENC values indicating increased codon usage bias.

The percentages of Bacteria, Archaea, Eukarya, and viruses making up the communities were not discernibly different between unfertilized and fertilized treatments (P =0.46, 0.37, 0.39, 0.21, respectively). In Bacteria and Eukarya, the percentages varied relatively little between samples within and across treatments (R^2^ = 9.6% and 13.5%, respectively). In Archaea and viruses, the mean percentages were substantially different between treatments (43 and 46% lower mean percentages in the fertilized treatment compared to the unfertilized treatment, respectively), but due to the high within-treatment variance, treatment explained limited variation in the relative abundances of these domains (R^2^ = 14% and 27%, respectively). The mean relative abundances in the unfertilized versus fertilized treatments were, respectively, 94% versus 93% in Bacteria, 6.0% versus 6.6% in Eukarya, 0.40% versus 0.23% in Archaea, and 0.05% versus 0.03% in viruses. Thus, Bacteria dominated the communities in both treatments. However, as expected, nutrient enrichment altered microbial community composition at a finer phylogenetic resolution, as indicated by the PCoA plot of community phylogenetic composition (based on the APIS analyses; Fig. S2) and supported by statistical analysis: the PcoA scores from a two-dimensional analysis differed between treatments (first dimension: R^2^ = 45.6%, P =0.096; second dimension: R^2^ = 51.3%, *P* = 0.070) and several taxonomic groups changed in abundance with fertilization (Fig. S3 and S4, all P < 0.01). Our results thus confirm the rarity of microbial Eukarya in the pond and show modest changes in the group’s relative abundance between treatments, indicating that the tRNA counts and our other bioinformatic analyses reflect mostly the response in bacteria.

Accompanying changes in fine-scale phylogenetic structure, we observed changes in several components of the genomic signatures of growth and trophic strategy (Fig. 1). Consistent with *a priori* predictions, mean estimated genome size of bacteria was 25% higher in the fertilized treatment (P =0.011), with nutrient enrichment explaining 75% of the variation between samples in mean genome size (Fig. 1). GC content of open reading frames was 9.9% higher in the fertilized treatment 54% compared to 49% in the unfertilized treatment (P =0.007, R^2^ = 86%; ENCp program).

Consistent with predictions, genomic features indicative of adaptations for maintaining high rates of transcription and translation were also positively associated with fertilization. The per sequence occurrence rate of tRNA genes, and the total number of tRNA genes per community were 93% and 64% higher, respectively, in the fertilized treatment (P < 0.001 and P =0.065 with R^2^ = 53%, respectively), and the residuals were also higher after accounting for the correlation between log number of tRNA genes and log total number of reads per sample to control for differences in sequencing depth (P =0.087, R^2^ = 52%). Likewise, the per sequence occurrence rate of 16S rRNA genes and total number of rRNA operons per community were 119% and 86% higher, respectively, in the fertilized treatment (P < 0.001 and P =0.096 with R^2^ = 50%, respectively), and the residuals after regressing log number of rRNA operons versus log total number of reads per sample were also significantly higher (P =0.038, R^2^= 52%). Fertilization explained 65% of the co-variation in these two traits (number of rRNA operons and tRNA genes) along a single axis, which was quantified by principal component analysis and provides a measure of the protein synthesis capacity of organisms (P = 0.031, unequal variances).

Consistent with *a priori* predictions, nutrient enrichment also increased codon usage bias in ribosomal protein genes (which are universally highly expressed genes) according to several measures of codon usage bias (Fig. 2). The mean effective number of codons (ENC) and ENC’ of a metagenome’s ribosomal protein sequences decreased with fertilization by 6.7 and 4.8%, respectively, indicating increased codon usage bias (ENC: P =0.018; R^2^ = 66%; ENC’: P =0.031; R^2^ = 55%). Median ENC and ENC’ were also lower in the fertilized treatment (P = 0.006, R^2^ = 75% and P =0.010; R^2^ = 75%) and Kolmogorov-Smirnoff tests indicated that the distributions in ENC and ENC’ values of ribosomal protein sequences significantly differed between treatments (all P <0.01, Fig. 2).

**Figure 2.**
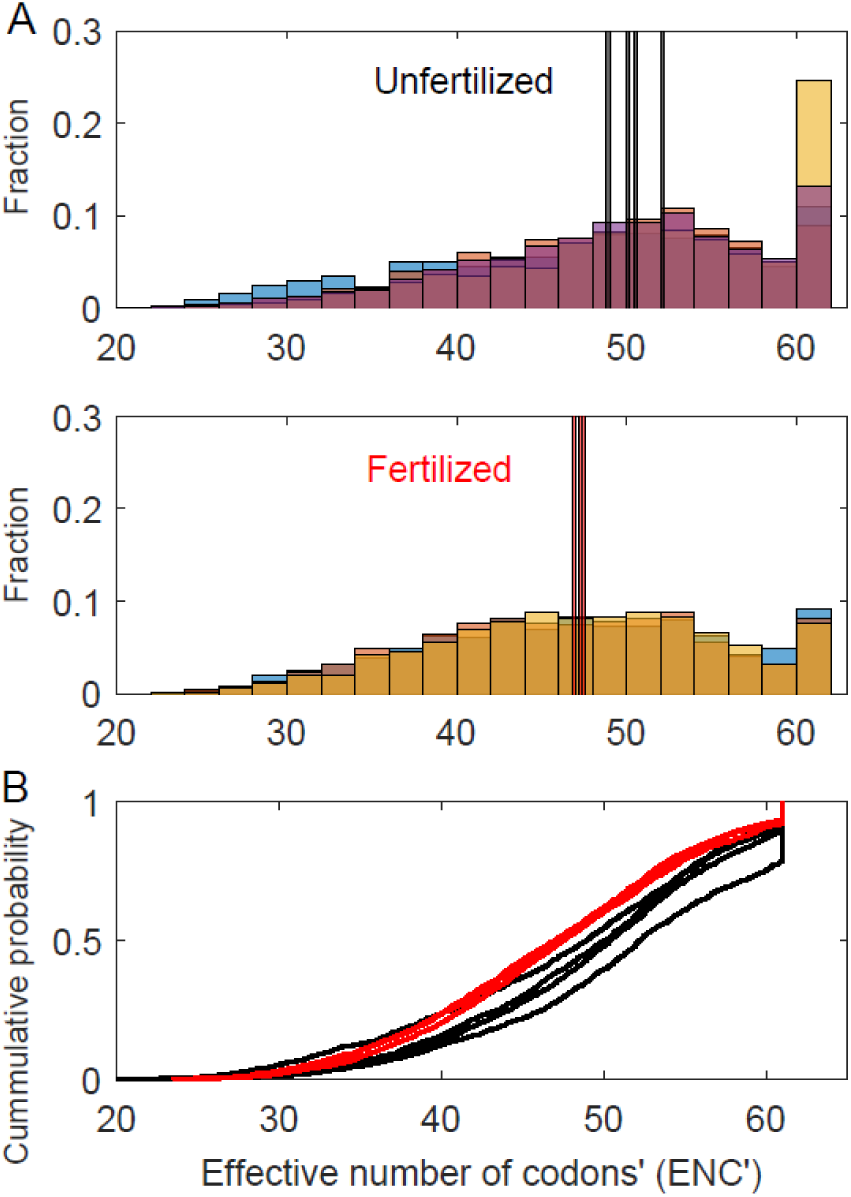
Histograms and cumulative distributions of the codon usage biases (ENC’ values) of the ribosomal protein sequences in the metagenomes, with lower ENC’ values indicating increased codon usage bias and thus increased speeds and/or accuracies of translation of ribosomes. (**A)** Tall vertical lines indicate the medians for each sample. Bar colors and their overlapping shades indicate different samples. **(B)** Y-axis shows the proportion of ENC’ values that are less than or equal to the ENC’ value designated by the plotted curves (fertilized = red curves, unfertilized = black curves).

As suggested by Figs. 1 and 2, we also observed that the variance in the P:C ratio and in mean and median codon usage bias of communities (quantified by ENC’, the more powerful indicator of codon usage bias) substantially decreased with fertilization by around an order of magnitude or more—by factors of 8, 31, and 27 (P:C ratio, P = 0.058; mean ENC’, P = 0.056, median ENC’, P =0.069; Table S1). Variance in genome size and mean and median ENC also decreased substantially by factors of 10, 10, and 6, respectively but these responses are less certain (genome size, P = 0.174; mean ENC: P = 0.169; median ENC: P = 0.275; Table S1). In contrast, variance in the log-transformed number of tRNA genes and rRNA operons substantially increased with fertilization, by 846% and 655% respectively (rRNA, P < 0.001; tRNA, P =0.013), while GC content variance did not appear to exhibit any change (P =0.56; Table S1).

Finally, we examined how well nutrient enrichment predicted the covariation of these genomic traits along a single principal component axis representing the oligotrophy-copiotrophy adaptation continuum. Principal components analysis revealed that nutrient treatment predicted 83% of the variation in genomic trait composition along this axis of trophic strategy (P =0.023; Fig. 3).

**Figure 3.**
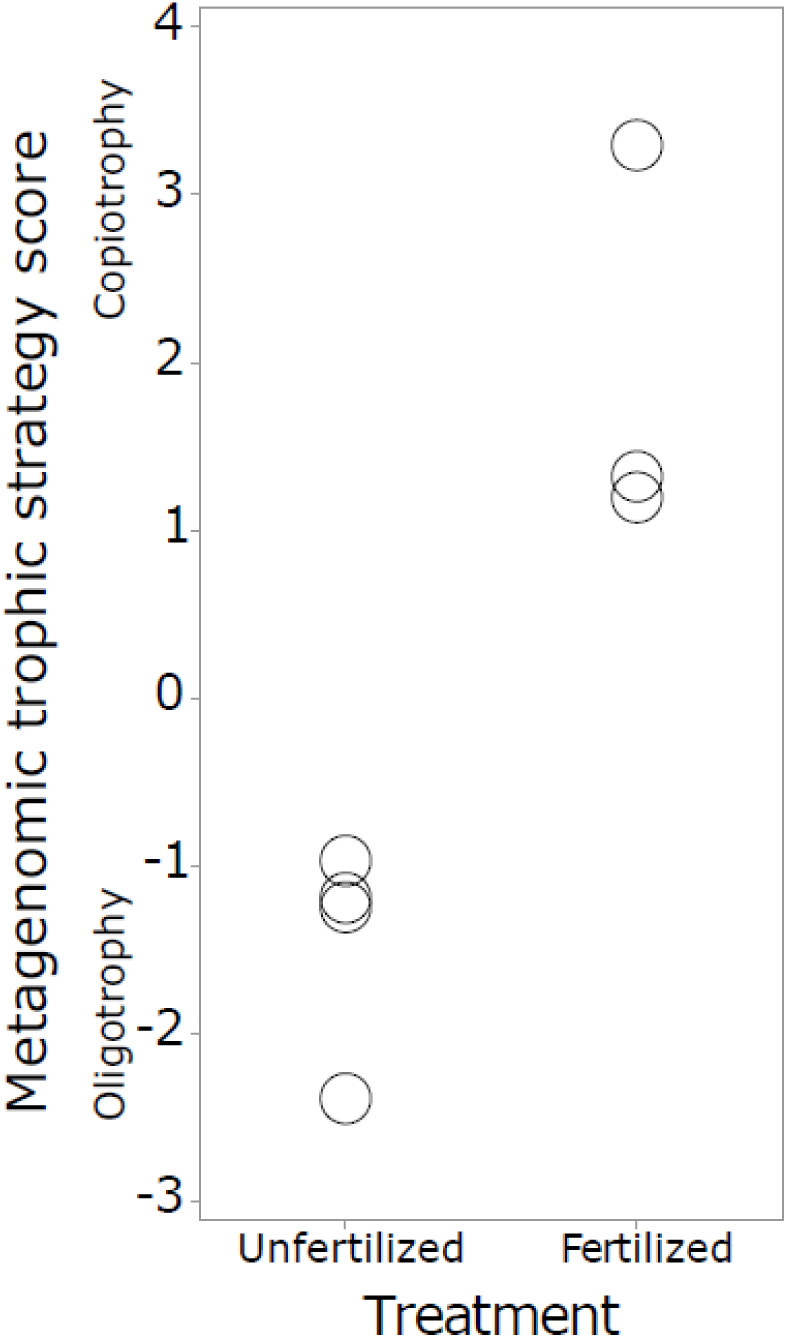
The principal component scores for the co-variation in community trait composition along a single dimension quantifying the oligotrophy-copiotrophy selection continuum for information processing traits. Higher scores indicate increased adaptation in information processing for copiotrophy (increased maximum growth rate in resource abundant environments), whereas low scores indicate increased selection for oligotrophy (increased resource use efficiency in low nutrient environments). Nutrient enrichment predicts 83% of the variance of genomic trait composition along this axis of growth and trophic strategy (R^2^ = 83%, P =0.004).

## Discussion

Our fertilization study demonstrated strong nutrient limitation in the Lagunita ecosystem, as manifested by increased biomass, chlorophyll content, and P:C ratios of suspended biomass in the fertilized pond versus the unenriched mesocosms. Using metagenomics, we report strong differences in microbial traits between fertilized and unfertilized treatments. We interpret these trait differences as reflecting differences in the growth and nutrient conditions of the two treatments rather than the difference in volume of the treatments, since the trait differences agree with all five of our directional predictions and there are no obvious reasons or existing theoretical work to indicate that the difference in habitat size between treatments should lead to the observed trait differences. Conservationists and ecologists should have an involved discourse in order to deliberate as to whether it is worth conducting future research to verify this interpretation by performing much larger experiments across multiple ponds (with wider environmental impacts). We also interpret these differences primarily as reflecting ecological dynamics —the differential success of lineages present within the pond and colonization of other lineages—rather than microevolutionary changes within populations, because the relative short time period of our experiment (32 days) encompasses a relatively low maximum number of generations of replication (SI) and so limited opportunity for measurable phenotypic evolution (e.g., Lenski and Travisano, 1994).

Overall, our study suggests that ecologically-significant phenotypic information about communities can be uncovered using a read-based approach (metagenomics) that leverages the relevant abundance and DNA characteristics of conserved genetic elements. Importantly, this approach avoids the biases associated with the massive gaps present in microbial taxonomic databases and cross-taxonomy assembly efficiencies (Kunin et al., 2010; Sogin et al., 2006; Temperton and Giovannoni, 2012; Yooseph et al., 2010).

### An oligotrophy-copiotrophy selection gradient in information-processing traits

Our genomic data (Figures 1-2) provide experimental evidence supporting all five of the directional predictions discussed in the introduction. The congruence of all trait responses with these predictions suggests a role for information-processing traits in community assembly and the geographic distribution of taxa. Compared to the unenriched reference systems, in the enriched pond we saw increased codon usage bias towards more efficient and precise translation (Fig. 1E and 2). We also saw an increase in the number of rRNA and tRNA genes, reflecting increased abundances of taxa with increased translational capacity for rapid growth. Such a result, in conjunction with the observed increase in the biomass P:C ratio, supports the “Growth Rate Hypothesis”—that fast-growing cells should have higher P-contents due to requiring higher concentrations of P-rich ribosomal RNA (Elser et al., 2000). Mean genome size was higher in the fertilized pond, reflecting the need to encode the larger array of genes needed to support production of a large translational and catabolic capacity in faster growing cells and the need to maintain a streamlined genome requiring minimal resources for maintenance and replication in nutrient-poor conditions (Fig. 1B). GC content also increased under fertilization, consistent with reduced relative abundances of oligotrophic taxa that maintain low GC-content as a resource conservation strategy (Fig. 1F).

Nutrient enrichment explained 50% or more of the variation in each trait and 88% of the co-variation of these five traits along a single statistical dimension that quantifies the oligotrophy-copiotrophy adaptation continuum (Fig. 3). In contrast with Vieira-Silva and Rocha (2010), who found no significant interspecific correlation of the traits GC content and genome size with generation time (Vieira-Silva et al., 2010), we found that GC content and genome size changed with nutrient enrichment, as well as with the other traits. The difference in results may reflect the presence of confounding variables and pronounced statistical error (measurement and biological) inherent to interspecific microbial growth and trait databases that are collated from a variety of sources, as in their study, highlighting the value of conducting *in situ* experiments.

Overall, however, our findings are largely consistent with observational studies of metagenomes and genomes along productivity gradients and comparisons across species varying in growth rate (Allen et al., 2012; DeLong et al., 2010; e.g., Foerstner et al., 2005; Lauro et al., 2009; Raes et al., 2007; Roller et al., 2013; Swan et al., 2013). Our experiment on communities *in situ* suggests that much of the variation in these genomic traits along productivity gradients or across species can similarly be attributed to the effects of the oligotrophy-copiotrophy strategy continuum and possibly related selection continua and tradeoffs in adaptive strategies, such as *r/K* selection and Grime’s “C-S-R Triangle” (e.g., Grime and Pierce, 2012) with *r*-selected species being more dominant in high-nutrient ecosystems.

### Variance changes in the genomic traits

We also observed changes in the variances in several community level traits (Table 1). As the direction of these changes differed among the traits in response to enrichment, the variance responses do not simply reflect potential for increased dispersal limitation between unfertilized mesocosms compared to enriched pond samples (see *Methods*). Instead, all-else-being equal, they suggest the strength of a trait’s role in mediating community assembly (i.e., in filtering the community) is different between unenriched and enriched treatments. The significant reduction in variance levels of P:C ratio and codon usage bias in response to fertilization suggest these traits play a stronger role in community assembly in the fertilized treatment, as discussed earlier. In contrast, the increased variance in rRNA operon and tRNA gene numbers in response to fertilization suggests a stronger, more consistent community assembly role for these traits in oligotrophic conditions. The augmented P and N requirements of taxa having high rRNA and tRNA gene copy numbers, which tend to maintain larger pools of ribosomes and other transcription machinery, may be particularly detrimental in oligotrophic conditions (Stevenson and Schmidt, 2004). In nutrient-rich conditions, on the other hand, some organisms may still do well even with fairly low numbers of these genes, for example: by having multiple genome copies; by mechanisms involving increasing concentrations and improved kinetics of RNA polymerase, rRNA and tRNA; or by having specialized niches that support only low growth rates, regardless of an organism’s rRNA operon copy number or the bulk phosphorus status of the ecosystem.

In contrast to shifts in variance observed for these traits, we observed no change in variance in GC content between treatments. Apparently, there is environmental filtering for low GC content in oligotrophic conditions, as expected given the higher nitrogen requirements and energetic expense of high GC content (Bragg and Hyder, 2004; Rocha and Danchin, 2002). However, it is less clear why there should be filtering for high GC content in response to nutrient enrichment, pointing to a need for further investigations into the implications of GC content for microbial physiological and community ecology.

There was no statistically significant change in variance in genome size, although we observed a suggestive 10-fold decrease in variance, suggesting that, if anything, genome size has a more variable ecological role in oligotrophic ecosystems. Based on genome streamlining theory alone, which suggests small genomes should be favored in oligotrophic conditions due to the favoring of cellular architectures that minimize resource requirements (Giovannoni et al., 2014; Morris et al., 2012), we would expect reduced variance in genome size in oligotrophic environments, in contradiction with our results. Consideration of metabolic scaling theory and co-evolution may provide the explanation. In natural microbial communities, including Cuatro Cienegas basin communities, many genomes rely on public goods and mutualisms (Embree et al., 2015; Rodríguez-Torres et al., 2017; Xavier, 2011). This co-dependence may allow some medium-sized genomes to persist in oligotrophic environments. In contrast, since there is presumably an interspecific increase in active mass-specific metabolic rate and r_max_ with genome size in bacteria (DeLong et al., 2010), under abruptly enriched conditions, members having larger genomes may displace the complex and diverse original community by growing faster.

### Concluding remarks

The genomic traits studied here all affect the costs and rates of information processing within cells and all of these traits responded as predicted to nutrient enrichment. Thus, optimizing the efficiency and capacity of cellular information processing appears to play a vital role in the evolutionary specialization of a microbe’s cellular biology to the particular trophic conditions of an ecosystem. The importance of such efficiency and capacity is even evident at the level of genes and protein sequences, as in the fertilized treatment we observed a notable increase in the number of ribosomal protein sequences with extremely high codon usage bias (ENC < ∼37) and a decrease in the number of ribosome protein sequences having nearly no codon usage bias (ENC > ∼60, Fig. 2). Information processing traits should thus be included in the development of trait-based theories and frameworks for microbial community ecology.

By applying a trait-based framework to metagenomics data from a replicated field experiment, we uncovered universal genomic traits that play an important role in governing the response of communities to altered growth and trophic conditions. We found that cells that dominate under nutrient-enriched conditions have increased capacities for information processing, whereas those found in oligotrophic conditions had genomic traits with reduced costs for information processing. Thus, all three core components of metabolism—information, energy, and material requirements and transformations—must be closely fine-tuned to the growth and trophic strategy of a microorganism.

## Acknowledgements

This study was conducted with financial support from NSF (DEB-0950175) and NASA (NAI5-0018) grants awarded to JJE, WWF-FCS to VS, and NSF (1536546) and NASA NAI (NNH05ZDA001C, NNH12ZDA002C, NNA08CN87A, NNA13AA93A) to JLS. This study was made possible with the sampling permit from Vida Silvestre-SEMARNAT, granted to V. Souza (09762). The WWF-FCS Alliance provided funds for VS and LEE. We thank J. Learned, J. Corman, J. Ramos, and E. Moody for field assistance, the Cuatro Cienegas community for its hospitality, and S. Vieira-Silva for advice and sharing of the codon usage bias analysis program *growthpred-v1.07.* This paper was written during a sabbatical leave of LEE and VSS in the University of Minnesota in Peter Tiffin and Michael Travisano laboratories, respectively, with support by scholarships from PASPA, DGAPA, UNAM.

## Accession Codes

Raw sequence data and metadata have been submitted to NCBI Sequence Read Archives, accessible through BioProject PRJEB22811.

## Supplemental Information

### Box S1. Further background on the genomic traits

#### Copy number of highly expressed genes essential to biosynthesis

In order to grow quickly, organisms must be able to transcribe genes, translate mRNA, and synthesize proteins at a sufficiently fast rate. Since ribosomes catalyze protein synthesis, increased capacity to replenish diminished ribosomal pools or maintain large standing stocks facilitates growth. Thus, faster growing organisms should tend to increase the number of rRNA operon copies (and so the 16S rRNA gene) in order to increase the rate of transcription of rRNAs, allowing the size and rate of turnover of the ribosomal pool to be larger and thereby preventing translation by the ribosomes from being the rate-limiting step in cellular growth.

Indeed, in bacteria and eukaryotes there are intraspecific and interspecific correlations between rRNA operon copy number per genome, which can vary by over an order of magnitude in bacteria (Acinas et al., 2004; Větrovský and Baldrian, 2013), and growth rate or generation time (Condon et al., 1995; Gyorfy et al., 2015; Klappenbach et al., 2000; Lauro et al., 2009; Shrestha et al., 2007; Stevenson and Schmidt, 2004; Yano et al., 2013). Thus microbes that are competitive in fast growing environments are likely to have to have a greater number of copies of the 16S rRNA gene.

Furthermore, a consequence of faster growing organisms having to have larger cellular quotas of rRNA is that they should tend to have higher phosphorus contents (Elser et al., 2003, 1996; Makino et al., 2003), since rRNA is phosphorus rich, and thus require more phosphorus-rich resources in order to build these rRNA pools. These observations, known as the “growth rate hypothesis” (Elser et al., 2000), have received considerable empirical support from comparative studies, but limited research has explored their usefulness as a predictive trait for community ecology.

Likewise, since the concentrations of tRNAs affects the rate of protein translation, having more copies of tRNA genes, which, for example, vary from under 30 to over 120 per genome in bacteria, may help a cell maintain larger pools of tRNAs, as has been observed (Higgs and Ran, 2008; Kanaya et al., 1999) and thereby maintain the faster translation rates required for faster growth rates. Indeed, number of tRNA genes has been shown to be negatively correlated with generation time (Higgs and Ran, 2008; Rocha, 2004).

#### Genome size

Genome size is a complex trait affecting multiple aspects of an organism’s ecology, physiology, and molecular biology, making it both an important trait to study as well as a challenging one since its influence on ecology may differ between environments, organisms, and historical circumstances. Although previously many microbiologists thought there was no relationship between growth rate and genome size (Mira et al., 2001), recent work suggests that among species growth rate and genome size may be positively correlated (DeLong et al., 2010), although this remains controversial (Vieira-Silva et al., 2010; Vieira-Silva and Rocha, 2010).

Additionally, bacteria in oligotrophic zones of the oceans have been reported to tend to have small genomes than the bacteria in copiotrophic zones (Allen et al., 2012; Lauro et al., 2009). Thus, organisms with larger genomes may, all else being equal, do better in high growth conditions and so respond favorably to nutrient fertilization.

There are several non-mutually exclusive reasons why genome size may be correlated with growth rate and response to fertilization. For one, because DNA is P-rich, large genome-sized organisms require more phosphorus to maintain and replicate their genomes and so they have greater difficulty than small genome size species in obtaining sufficient amounts of P in oligotrophic, P-limited environments. Larger genome-species also tend to have to have larger cells, leading to, on average, decreased surface-to-area-volume ratios, which in turn puts larger cells at a disadvantage for obtaining sufficient nutrients (Okie, 2013). Genome size also affects the size, structure and function of metabolic networks of organisms by allowing for a greater diversity of enzymes, a higher diversity of metabolic pathways, and enhanced metabolic multi-functionality, which may affect an organism’s ability to take up a variety of substrates, the speed of resource uptake and transformation, and the yield of resource transformation (DeLong et al., 2010; Maslov et al., 2009). Finally, larger genomes allow organisms to encompass greater copy numbers of highly expressed genes and genes involved in protein translation machinery, such as tRNA and rRNA genes, which in turn allow for greater translation rates, as discussed above.

#### Nucleotide base composition of DNA

The percentage of a genome’s DNA composed of the nucleotide bases guanine and cytosine is known as a genome’s GC content. Genome GC content varies from widely across life, and there appears to be pervasive selection on the GC content of bacterial genomes, leading to selection for high GC content in some bacterial genomes (Hildebrand et al., 2010). The reasons for positive selection on GC content are contested, and it is likely that there are multiple different selective forces on GC content related to the niches and environmental conditions of different species. Because GC bonds have eight nitrogen atoms whereas AT bonds contain only seven, higher GC content genomes tend to have higher nitrogen content (Bragg and Hyder, 2004), making higher GC content organisms have increased nutrient requirements for replication and DNA repair.

Rocha and Danchin (Rocha and Danchin, 2002) found that parasitic and symbiotic bacteria have lower GC content than free-living bacteria and propose an explanation based on the biochemical details of nucleotide metabolism and differences in the energetic expense of GTP and CTP nucleotides versus ATP and UTP nucleotides. The explanation boils down to the difference resulting from increased selection by competition for scarce resources in parasites and symbionts. Here we extend this explanation to include extremely oligotrophic genomes as similarly susceptible to these evolutionary forces as parasites and symbionts, and thus propose that an emergent consequence of these effects should be a positive association between GC content and growth rate. Suggestive support for this possibility is provided by the observation that GC content tends to be higher in DNA of environmental samples from complex environments, such as soils, and environments with high amount of nutrients, such as whale carcasses and copiotrophic waters, compared to simple and lower productivity environments such as oligotrophic pelagic communities in the Sargasso Sea (Allen et al., 2012; Foerstner et al., 2005; Raes et al., 2007).

#### Codon usage bias

An amino acid can be encoded by multiple different codons (nucleotide triplets), but these synonymous codons have different kinetic properties, including different probabilities of mistranslation. In highly expressed genes essential to growth, such as ribosomal protein genes, there should be increased selection for biasing the usage of certain synonymous codons over others in order to optimize the accuracy and speed of translation, especially in organisms with fast growth rates.

Since 18 of the 20 standard amino acids are encoded by two or more synonymous codons, genomes and genes may favor certain codons over others without altering translation products.

Different synonymous codons have different probabilities of mistranslation. Thus, the use of more accurately translated codons can be beneficial because mistranslation wastes energy, reduces the translation rate (all else being equal), and/or can cause cytotoxic protein misfolding. Codon usage bias (CUB) has been documented in bacteria, archaea, and eukaryotes (Satapathy et al., 2014; Subramanian, 2008; Vieira-Silva and Rocha, 2010). The highly expressed genes of a genome, such as ribosomal protein genes, tend to have greater CUB (Subramanian, 2008; Vieira-Silva and Rocha, 2010).). As CUB can alter the accuracy and speed of translation, this greater CUB presumably reflects selection for a greater translational accuracy and/or or efficiency (Hershberg and Petrov, 2008) which should increase rates of translation.

It follows from this positive effect of CUB on translation that faster-growing organisms should tend to have greater CUB, especially in genes that are highly expressed and essential for growth (such as ribosomal protein genes). Indeed, a negative relationship between an organism’s generation time and the CUB of its ribosomal protein genes has been reported (Subramanian, 2008; Vieira-Silva and Rocha, 2010). A few studies have compared the CUB of metagenomes from different environments (Roller et al., 2013; Vieira-Silva and Rocha, 2010), but more work is required to clarify the role of CUB in community assembly.

## Detailed methods

### Study site description

The Cuatro Ciénegas basin is an enclosed evaporitic valley in the Chihuahuan desert, Mexico. Despite its aridity, the CCB harbors a variety of groundwater-fed springs, streams, and pools. Past research has also shown that these aquatic environments are heavily mineralized (carbonate and sulfate) and stoichiometrically imbalanced, with high nitrogen (N):phosphorus (P) ratios indicative of strong ecosystem P limitation (Corman et al., 2016).

The whole-ecosystem fertilization experiment took place in Lagunita, a shallow pond adjacent to a larger lagoon (Laguna Intermedia) in the Churince flow system. Consistent with Churince’s geochemistry, Lagunita waters are high in conductivity and dominated by Ca^2+^, SO_4_^2-^, and CO ^2-^ (Lee et al., 2015, 2017). Evaporation rates are high and Lagunita’s volume is strongly reduced during summer. During the experiment, maximum water depth in Lagunita began at 32 cm and declined to 12 cm by the experiment’s end. The pond’s N:P stoichiometry is highly imbalanced, with an average molar TN:TP ratio of 122, indicative of strong P limitation, as previously demonstrated in this system during a mesocosm experiment completed in 2011 (Lee et al., 2015, 2017).

### Fertilization regime

Prior to initiation of fertilization, five replicate enclosures were established in different parts of the pond; these served as the unenriched, reference treatment. As in Lee *et al*. (2015, 2017), each enclosure consisted of a clear plastic tube 40 cm in diameter pushed ∼20 cm into the sediment and extending ∼20 cm above the water surface. A small number of fish and larger aquatic macroinvertebrates (∼1 cm or greater) were removed from the unfertilized mesocosms with a dip net before beginning the experiment. Such removals were necessary to ensure that enclosures did not experience unduly large stochastic disruption from animal activities resulting from differing numbers of large animals being trapped inside at unnaturally high densities. This would have confounded the difference in nutrient conditions between the enriched and unenriched treatments, undermining the purpose of the experiment. To the extent that such consumers augmented nutrient availability outside of the unenriched enclosures, then their removal from the enclosures would have accentuated the nutrient enrichment contrast between the fertilized pond and the unenriched enclosures, amplifying our planned experimental contrast.

The fertilization procedure was based on a previous mesocosm experiment in Lagunita that produced major shifts in microbial biomass and species composition in the water column (Lee et al., 2015, 2017). Prior to sampling, a morphometric map of the pond was created, allowing us to estimate the pond’s water volume and adjust that volume estimate as water depth changed. Based on the pond’s volume, we fertilized to increase PO_4_ concentration in the water by 1 uM (as KH_2_PO_4_).

As in one of the treatments in Lee et al. (2015), we also added NH_4_NO_3_ in a 16:1 (molar) N:P ratio with the added P. The soluble reactive phosphorus (SRP) concentration of the pond was then measured every 3-4 days (see *Routine water chemistry*), after which we added sufficient KH_2_PO_4_ to bring the pond’s *in situ* concentration back to 1 uM, along with the appropriate amount of NH_4_NO_3_ to achieve a 16:1 molar ratio. Fertilizer was added by mixing fertilizer solution with ∼2 L pond water and broadcasting the mixture into all regions of the pond.

### Field monitoring and sampling

Following initiation of fertilization, the pond and internal unfertilized mesocosms were sampled every 4 days to monitor basic biogeochemical and ecological responses. Water was sampled adjacent to and within each of the five internal mesocosms. Water samples were obtained by plunging a clean 2-L polycarbonate beaker ∼10-cm beneath the surface. Samples were filtered onto Whatman GF/C filters for analysis of chlorophyll (chl *a*) concentrations and passed through 0.2-µm polyethersulfone membrane filters (Pall Life Sciences, Port Washington, NY) and filtrate was used for analyses of nitrate (NO_3_), ammonia/ammonium (NH_3/4_), soluble reactive phosphorus (SRP), and total dissolved phosphorus (TDP). After 16 and 32 days of fertilization, water samples were also filtered onto pre-combusted Whatman GF/F filters for analysis of concentrations of C, N, and P in suspended particulate matter (seston). Unfiltered water samples were frozen for later analysis of total N (TN) and total P (TP) concentrations.

At the end of the 32-day experimental period, a set of samples was taken for metagenomic analysis; this involved five water samples from the pond itself (fertilized treatment) and one water sample from inside each of the five unfertilized internal mesocosms. Microplankton, including bacteria, in the water samples were filtered onto sterile GF/F filters (0.7-µm nominal pore size, Whatman, Piscataway, NJ, USA) and then frozen immediately in liquid nitrogen in the field and held at low temperature (<-80° C) until laboratory DNA extraction, purification, and sequencing. Note that given the 0.70 µm pore size, extremely small prokaryotes were not part of our metagenomes and so our results do not apply to these picobacterioplankton. If anything, their inclusion would augment predicted community-level trait responses to fertilization, since picoplankton are slow-growers, tend to do poorly in nutrient rich waters, have small genomes (Box 1) and so likely decrease in abundance in the fertilized treatment.

### Routine water chemistry

Chl *a* on GF/C filters was quantified fluorometrically using a TD-700 fluorometer (Turner Designs, Sunnyvale, CA) after 16–24 hours of extraction in cold absolute methanol (Arar and Collins, 1997). TN concentrations were measured using a Shimadzu TOC/TN analyzer. TDP and TP concentrations were measured using the same colorimetric method after persulfate digestion of the filtered or unfiltered samples, respectively. GF/F filters with seston were thawed, dried at 60°C and then packed into tin discs (Elemental Microanalysis, U.K.) for C and N analyses with a Perkin Elmer PE 2400 CHN Analyzer at the Arizona State University Goldwater Environmental Laboratory. Another set of dried GF/F filters prepared from the same water samples was used for estimating seston P content. These filters were digested in persulfate followed by colorimetric analysis for P.

### DNA extraction and sequencing

DNA was extracted from the GF/F filters with a slight modification (increasing volume of PW1 solution to 1.5 mL) using the MO BIO PowerWater DNA Isolation kit (MOBIO Laboratories, Inc., Carlsbad, CA, USA). Following extraction, DNA yield and quality were assessed by PicoGreen assay (Life Technologies, Carlsbad, CA, USA) and prepared for sequencing on Illumina MiSeq with 12 samples per v2 2x250bp sequencing run.

### Bioinformatics and data analyses

#### Annotation

Raw reads were trimmed of barcodes, quality filtered, and rarefied to 100,000 sequences per sample. For the quality filtering, we used the standard Qscore of 25. Two samples from the fertilized treatment and one sample from the unfertilized treatment were left out of subsequent analyses because they had sequencing depths less than 20% of the rest of the samples (whose sequencing depth averaged 2.5 x 10^6^ reads) and low quality scores. Sequences were phylogenetically annotated using the Automated Phylogenetic Inference System (APIS), which has hard-coded parameters and is designed to optimize annotation accuracy (Allen et al., 2012). Annotations from APIS were used to generate a Bray-Curtis distance matrix visualized with PCoA using the R package vegan (Oksanen et al., 2016).

#### Number of ribosomal RNA operons and tRNA genes

We used Phylosift (Darling et al., 2014) to annotate non-sub-sampled libraries, allowing for the counting of the number of 16S rRNA bacteria and archaea genes in each sample, which averaged 2607. Counting 16S rRNA genes provides a means of evaluating the number of rRNA operons, because in prokaryotes 16S RNA genes are typically transcribed as part of a rRNA operon and furthermore the typical situation in the genomes of cultivated organisms is that each bacterial rRNA operon has a single 16S rRNA gene (Grigoriev et al., 2011). The program tRNAscan-SE v1.4 (Lowe and Eddy, 1997), which is specifically designed for recognizing tRNA genes, was used with a general tRNA model in order to count the number of tRNA genes in non-sub-sampled libraries. In the *Statistical analyse*s section we describe our method for ensuring that results were not sensitive to variation in DNA sequencing depth. Although the tRNAscan-SE approach employed did not distinguish between Bacteria, Archaea and Eukarya, the observed rarity of Eukarya and insignificant changes in the group’s relative abundance between treatments (see *Results*) indicate that the tRNAscan-SE results, and our other bioinformatic analyses reflect variation in prokaryotes, namely Bacteria (Archaea are very rare in CCB samples, Lee et al., 2017), rather than Eukarya.

#### Genome size estimates

Bacterial genome sizes were estimated according to methods in Allen et al. (2012). Briefly, length normalized core marker gene counts identified as bacterial by APIS were used to determine number of genome equivalents in a sample. Total number of predicted proteins annotated as bacterial by APIS was then divided by number of genome equivalents.

#### Codon usage bias metrics (ENC and ENC’)

The average number of ribosomal protein sequences identified per sample was 2607. We estimated two commonly used metrics, the effective number of codons (ENC) (Wright, 1990) and a related measure, ENC’ (Novembre, 2002), to quantify the degree of codon usage bias of ribosomal protein gene sequences. Ribosomal protein genes are some of the most highly expressed genes, and so should experience augmented selection for codon usage bias, enhancing its translation, especially under favorable growth conditions (Vieira-Silva and Rocha, 2010). Both metrics are inversely related to the extent of codon usage bias and were calculated for each ribosomal protein gene sequence. ENC is relatively statistically well-behaved and insensitive to short gene lengths compared to other measures of codon usage bias. Its value can have a maximum of 61, representing the situation in which all codons are being used with equal frequency; smaller values indicate decreasing uniformity in codon usage within a sequence.

ENC’ has similar advantages as ENC but additionally accounts for departures in background nucleotide composition from a uniform distribution (Novembre, 2002).

In order to calculate the ENC and ENC’ values of ribosomal protein sequences, we first used Phylosfit (Darling et al., 2014) to identify the ribosomal protein gene sequences of Bacteria and Archaea and then used the program ENCprime (Novembre, 2002) to calculate ENC and ENC’ values for each identified sequence. For each sample, background nucleotide composition was estimated to be the average nucleotide frequencies of all the samples’ reads.

#### Statistical analyses

In order to evaluate differences between treatments in phylogenetic composition and in means of community-level traits, we used two sample T-tests conservatively assuming unequal variances (for traits, these tests were one-tailed with the alternative hypothesis based on the predicted differences between means in Box 1). We also used General Linear Models with treatment as a fixed factor in order to test for the effects of fertilization on the codon usage bias of individual ribosomal protein sequences and to determine the amount of variation (R^2^) in traits explained by the treatment. Numbers of tRNA genes and rRNA operons were log-transformed before these analyses to improve normality. We also employed single-tailed Poisson rate tests to examine differences in the numbers of rRNA operons and tRNA genes, as the Poisson rate test provides a more powerful test than the T-test for examining differences in the per sequence rate of occurrences of tRNA genes and rRNA operons between treatments, where the sample size for each treatment is the number of DNA samples and the treatment’s observation length is the mean number of reads per metagenome.

In order to account for potential effects of sequencing depth on the number of rRNA operons and tRNA genes, we regressed the numbers against a sample’s log total number of reads and then performed GLM analyses on the residuals (ascertaining whether or not for a given sequencing depth samples from the fertilized treatment have higher numbers of these genes, that is, higher residuals). Two-sided Bonett’s tests were used to test for the effect of fertilization on variance between samples of the community-level traits. Kolmogorov-Smirnoff tests were conducted to evaluate whether or not within-sample distributions of ENC and ENC’ values of ribosomal protein gene sequences differ between treatments.

Finally, we examined how well nutrient enrichment predicted the covariation of these genomic traits along a single axis that quantified molecular adaptiveness to oligotrophic versus copiotrophic conditions. Principal components analysis (PCA) of each samples’ sets of community trait values was used to place each metagenome along a single principal component representing the studied genomic features. Before performing PCA, in order to give equal weight to each trait, variables were first standardized by subtracting means and dividing standard deviations.

#### Calculations on the potential for evolutionary change during the experiment

For microorganisms with 2-h minimum generation times—the approximate average minimum generation time of prokaryotic isolates in many data sets (e.g., DeLong et al., 2010; Vieira-Silva and Rocha, 2010)—a maximum of 384 generations of replication can occur in the 32-day period of the experiment, which is barely sufficient for much evolutionary change (Lenski and Travisano, 1994), especially of some of the genome-scale traits like GC content and codon usage bias. Many of the microorganisms in natural ecosystems (that is, the unculturable taxa underrepresented in generation time datasets and that make up the majority of prokaryotic communities) likely have minimum generation times greater than 2 hours and under field conditions do not achieve or sustain their minimum generation times for the whole course of the experiment. Thus we expect that in our experiment, populations experienced much fewer than 384 generations, providing very limited opportunity for evolutionary change to affect the genomic traits.

---

**Figure S1.**
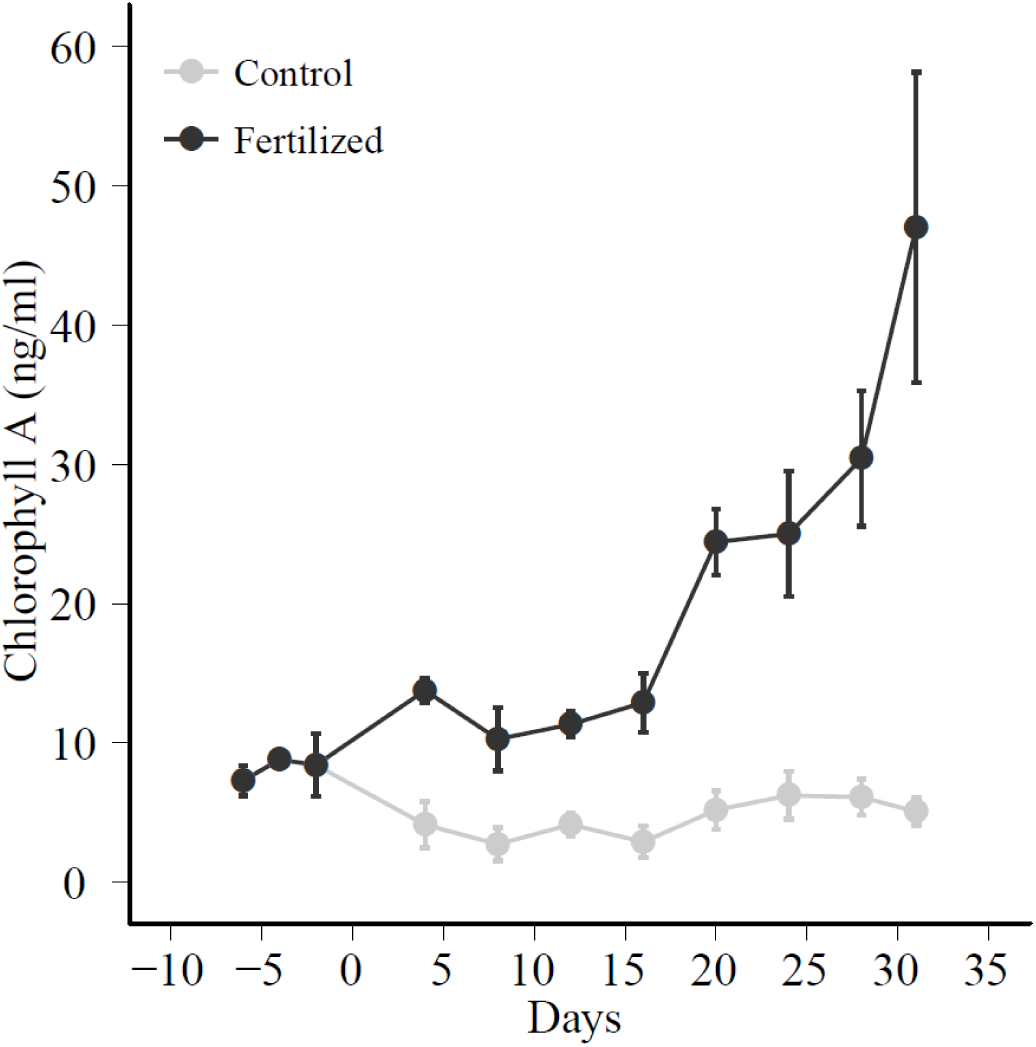
Chlorophyll *a* concentration increased with nutrient enrichment whereas it remained relatively invariant in the unfertilized treatment.

**Figure S2.**
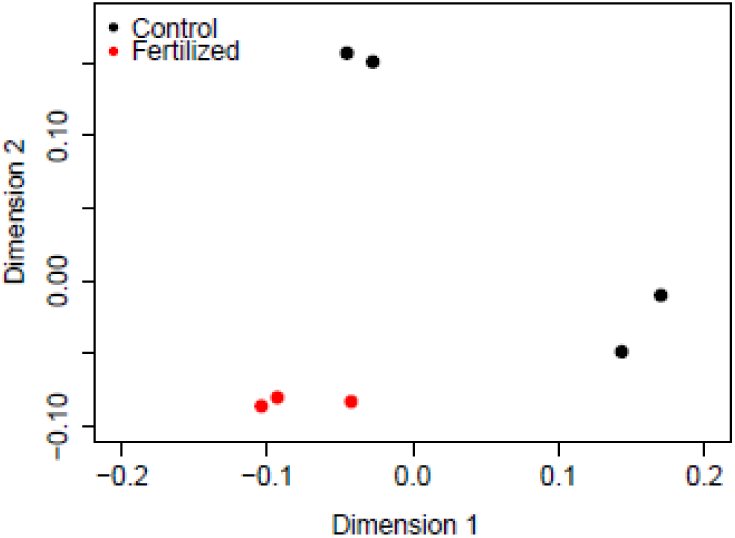
Principle coordinates analysis (PCoA) of the community phylogenetic structure inferred from the metagenomes shows that the phylogenetic composition of samples, as indicated by each point, is substantially different between unfertilized and fertilized treatments. Microbial community phylogenetic composition also varied notably within the unfertilized mesocosms, falling into two clusters driven by variation in the relative abundance of Alphaproteobacterum, whereas enriched communities all shared relatively similar phylogenetic composition, indicating a convergence of effects of fertilization on community composition.

**Figure S3.**
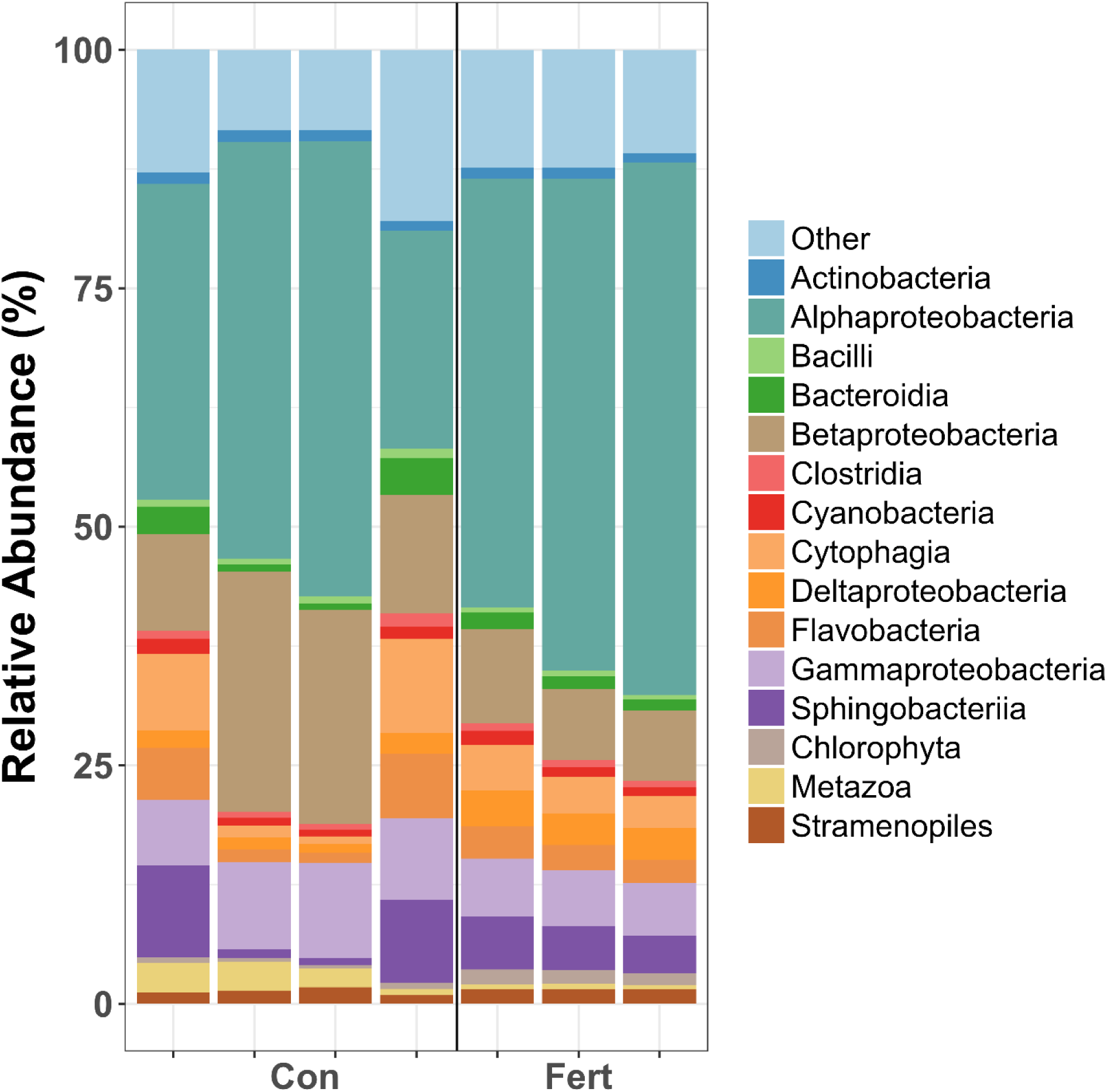
Relative abundances of taxonomic phyla in the samples from the unfertilized (Cont) and fertilized (Fert) treatments.

**Figure S4.**
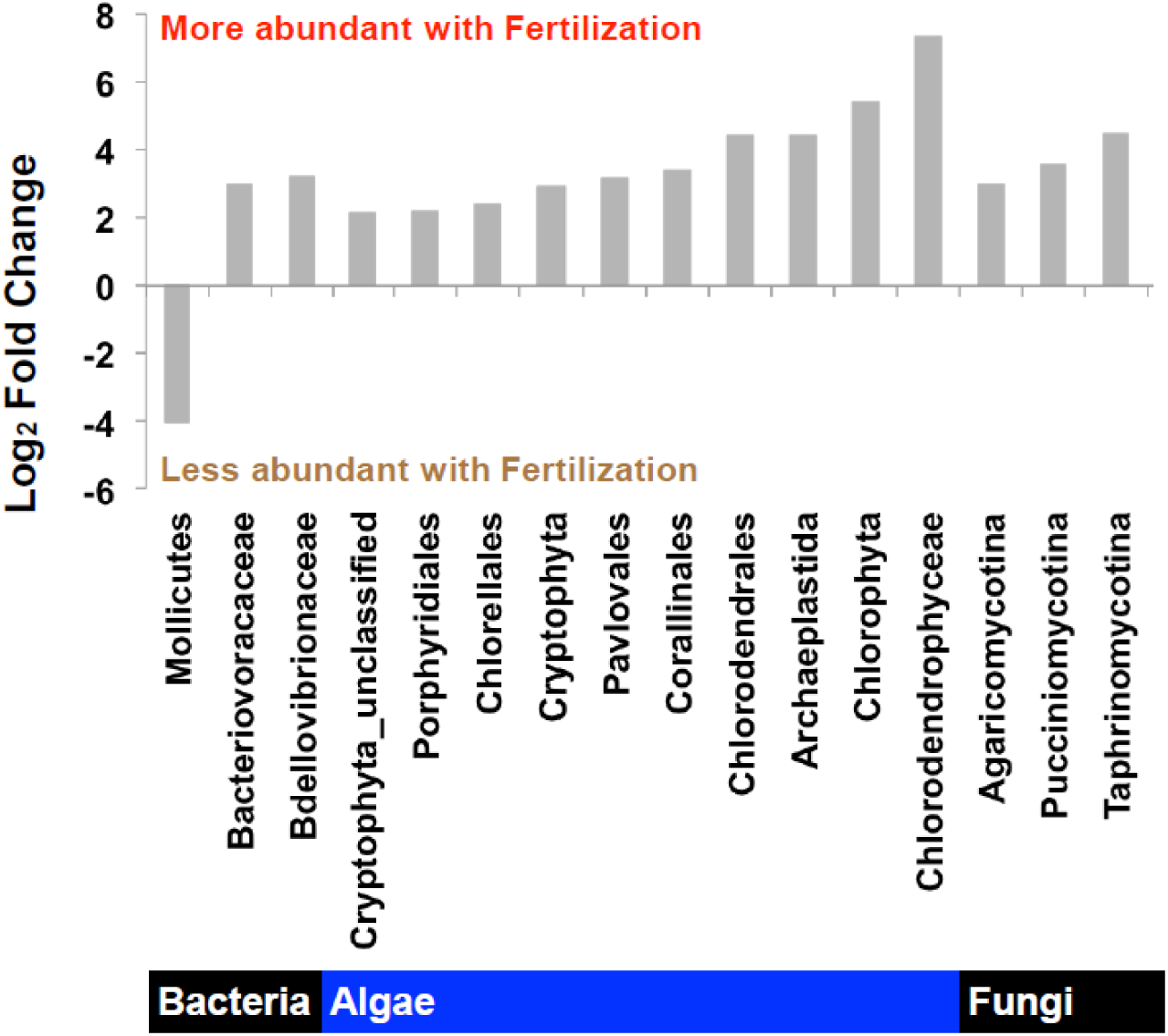
The significant relative abundance changes in taxonomic groups suggested by the statistical analysis of the phylogenetic composition determined by APIS. The statistical analysis is based on the statistical methods developed in the EdgeR package (Robinson et al., 2010). Note that less than 50% of reads could be annotated. All P < 0.01.

**Table S1.**
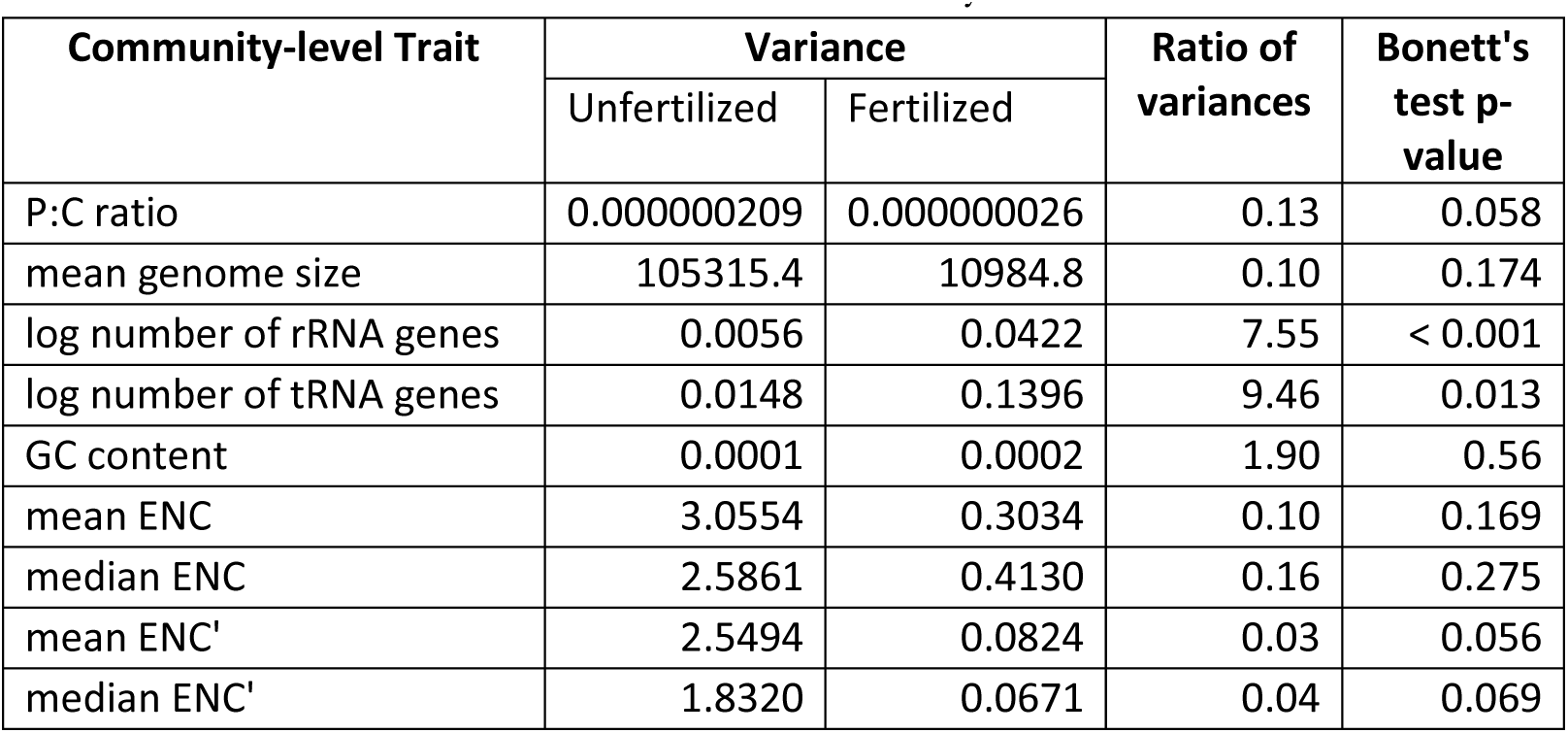
Differences in the variance of community-level traits between treatments.

